# Next-generation phylogeography resolves post-glacial colonization patterns in a widespread carnivore, the red fox (*Vulpes vulpes*), in Europe

**DOI:** 10.1101/2020.02.21.954966

**Authors:** Allan D. McDevitt, Ilaria Coscia, Samuel S. Browett, Aritz Ruiz-González, Mark J. Statham, Inka Ruczyńska, Liam Roberts, Joanna Stojak, Alain C. Frantz, Karin Norén, Erik O. Ågren, Jane Learmount, Mafalda Basto, Carlos Fernandes, Peter Stuart, David G. Tosh, Magda Sindicic, Tibor Andreanszky, Marja Isomursu, Marek Panek, Andrey Korolev, Innokentiy M. Okhlopkov, Alexander P. Saveljev, Boštjan Pokorny, Katarina Flajšman, Stephen W. R. Harrison, Vladimir Lobkov, Duško Ćirović, Jacinta Mullins, Cino Pertoldi, Ettore Randi, Benjamin N. Sacks, Rafał Kowalczyk, Jan M. Wójcik

## Abstract

Carnivores tend to exhibit a lack of (or less pronounced) genetic structure at continental scales in both a geographic and temporal sense and this can confound the identification of post-glacial colonization patterns in this group. In this study we used genome-wide data (using Genotyping-by-Sequencing (GBS)) to reconstruct the phylogeographic history of a widespread carnivore, the red fox (*Vulpes vulpes*), by investigating broad-scale patterns of genomic variation, differentiation and admixture amongst contemporary populations in Europe. Using 15,003 single nucleotide polymorphisms (SNPs) from 524 individuals allowed us to identify the importance of refugial regions for the red fox in terms of endemism (e.g. Iberia). In addition, we tested multiple post-glacial re-colonization scenarios of previously glaciated regions during the Last Glacial Maximum using an Approximate Bayesian Computation (ABC) approach that were unresolved from previous studies. This allowed us to identify the role of admixture from multiple source population post-Younger Dryas in the case of Scandinavia and ancient land-bridges in the colonization of the British Isles. A natural colonization of Ireland was deemed more likely than an ancient human-mediated introduction as has previously been proposed and potentially points to an increased mammalian fauna on the island in the early post-glacial period. Using genome-wide data has allowed us to tease apart broad-scale patterns of structure and diversity in a widespread carnivore in Europe that was not evident from using more limited marker sets and provides a foundation for next-generation phylogeographic studies in other non-model species.

## Introduction

Over the last 30 years, phylogeographic studies have highlighted the roles of major past climatic and geophysical events in shaping contemporary genetic structure and diversity in a multitude of species (Avise et al., 1987; Pedreschi et al., 2019; Stojak, Borowik, McDevitt, & Wójcik, 2019; Taberlet, Fumagalli, Wust-Saucy, & Cosson, 1998). During the Last Glacial Maximum (LGM; ~27-19 thousand years ago (kyrs BP; Clark et al., 2009)) many terrestrial plant and animal species retreated, and were often restricted, to refugial areas (Sommer & Nadachowski, 2006; Taberlet et al., 1998). In Europe, several overarching patterns have emerged from fossil data and phylogeographic studies, with refugia identified in the three ‘classic’ Mediterranean (Iberian, Apennine and Balkan) peninsulas (Hewitt, 1999; Sommer & Nadachowski, 2006; Taberlet et al., 1998), but also further north in areas in or adjacent to the Carpathians mountains and the Dordogne region in France as examples of ‘cryptic refugia’ (McDevitt et al., 2012; Provan & Bennett, 2008; Stojak et al., 2016). Phylogeographic studies of the most widely studied group, the terrestrial mammals, have shown distinct mitochondrial DNA (mtDNA) lineages in small mammals (Searle et al., 2009; Stojak et al., 2016; Vega et al., 2020) and ungulates (Carden et al., 2012; Sommer et al., 2008) that are consistent with contraction and re-expansion from these refugial regions (Hewitt, 1999; Taberlet et al., 1998).

Carnivores appear to be an exception to this general pattern however, with either a lack of, or less pronounced phylogeographic structure shown across continental scales (Frantz et al., 2014; Hofreiter et al., 2004; Korsten et al., 2009; Mucci et al., 2010). One such carnivore, the red fox (*Vulpes vulpes*), is well-represented in the fossil record in Europe (Sommer & Benecke, 2005) and has numerous records during the LGM in recognized refugial areas such as the Mediterranean peninsulas, and further north in areas in or adjacent to the Carpathian mountains, and the Dordogne region in France (Sommer & Benecke, 2005; Sommer & Nadachowski, 2006). Previous studies using mtDNA on modern and/or ancient specimens have revealed a general lack of genetic structure on a continental-wide scale, in both a geographic and temporal sense (Edwards et al., 2012; Kutschera et al., 2013; Teacher et al., 2011). The lack of phylogeographic structuring in the red fox and other carnivores has been previously attributed to these species persisting outside the traditional refugial areas during the LGM, and effectively existing as a large interbreeding population on a continental scale (Edwards et al., 2012; Teacher et al., 2011). However, despite the abundant fossil data for red foxes, there is a distinct lack of fossils from central Europe or further north during the LGM (Sommer & Benecke, 2004, 2005).

More recent studies (Statham et al., 2014, 2018) identified mtDNA haplotypes that were unique to particular regions (e.g. Iberia) that potentially indicate long-term separation from other European populations. The concerns about the use of short mtDNA sequences is that they may not fully capture the signals of post-glacial colonization patterns in species with high dispersal capabilities (Keis et al., 2013; Koblmüller et al., 2016). One solution is the utilisation of microsatellite markers in conjunction with mtDNA data (Statham et al., 2018). However, several carnivore species (e.g. badgers *Meles meles* and otters *Lutra lutra*) show a similar lack of resolution in terms of broad-scale genetic structure across continental Europe using microsatellites (Frantz et al., 2014; Mucci et al., 2010). For the red fox, several distinct groups in Europe were identified using microsatellite markers from Bayesian clustering analysis, with distinction between foxes in Ireland, Britain, Spain, Italy and Scandinavia being apparent (Statham et al., 2018). The rapid mutation rate of microsatellites leaves it unclear whether divisions reflect ancient isolation or more recent population structure, owing to recently limited gene flow. This uncertainty and lack of resolution in phylogeographic structure outside the ‘traditional’ (and well-established) refugia means that inferring post-glacial colonization patterns of previously glaciated regions during the LGM is challenging with more limited genetic marker sets. This is particularly evident in the British Isles and Scandinavia where these regions present differing problems in terms of how and when contemporary populations of terrestrial species reached these areas in post-glacial periods (Carden et al., 2012, 2020; Herman et al., 2014).

The island of Ireland has long presented a biogeographical quandary in terms of how and when terrestrial species colonized it (Carden et al., 2012, 2020; McDevitt et al., 2011). The area’s latitude means that it was covered almost entirely by the ice sheet during the LGM and it didn’t become an island until approximately 15,000 years ago (almost twice as long as Britain; Edwards & Brooks, 2008). Because of this, humans have been proposed as the primary mechanism of transport for its mammalian fauna in ancient and modern times (Carden et al., 2012; Frantz et al., 2014). An estimate of 10.2 kyrs BP was estimated for a split between Irish and British red foxes using mtDNA data, but with a 95% CI range that incorporated the possibility of natural colonization before Ireland became an island (Statham et al., 2018). Haplotype diversity at different mtDNA markers is high in Irish red foxes (Edwards et al., 2011; Statham et al., 2018) which would appear to contradict a more recent, human-mediated origin that has been inferred from the only fossil dated to the Bronze Age (3.8 kyrs BP; Sommer & Benecke, 2005). Indeed, natural colonization of other species (e.g. stoat *Mustela erminea*) have been proposed over post-glacial land bridges (Martínková, McDonald, & Searle, 2007) and a recent re-evaluation of fossil dates has now placed multiple mammalian species on the island close to the end of the LGM period (Carden et al., 2020). Therefore, a more thorough investigation of the red fox’s colonization history on the island has implications beyond this species only.

Fennoscandia (present-day Sweden, Norway, Finland and Denmark) was covered by a huge ice sheet during the LGM and would have been inhospitable to most species until its complete retreat approximately 10,000 years ago (Patton et al., 2017). However, some parts of southern Sweden and western Norway would have had ice free regions during the later Younger Dryas Glaciation (12.9–11.7 kyrs BP) and several temporary land-bridges connected these countries to Denmark between approximately 13.1 and 9.2 kyrs BP (Herman et al., 2014; Marková et al., 2020). Although glacial refugia in Scandinavia during the LGM have been proposed for several species, including some mammals (Lagerholm et al., 2014; Westergaard et al., 2019), the red fox does not appear in the fossil records in southern Scandinavia until after ~9,000 yrs BP (Sommer & Benecke, 2005). Based on evidence from mtDNA, microsatellites and Y chromosome data, multiple colonization events have been proposed from the south and east for the red fox (Norén et al., 2015; Wallén et al., 2018) but the progression of these events remain untested. Several mammalian species have been proposed to have colonized from the south over temporary land-bridges and persisted through the Younger Dryas before being supplemented with further colonization wave(s) from the east (Herman et al., 2014; Marková et al., 2020). It is unclear however if this is a frequently-used post-glacial colonization route for many of the mammals that are currently present in the region.

The advent of next-generation sequencing technologies holds great promise for phylogeographic studies, allowing for thousands of single nucleotide polymorphisms (SNPs) to be genotyped in non-model organisms and providing a representation of the organism’s entire genome (Garrick et al., 2015; McCormack, Hird, Zellmer, Carstens, & Brumfield, 2013). The use of reduced-representation techniques (e.g. genotyping-by-sequencing, GBS) has already demonstrated their potential in resolving phylogeographic patterns in non-model organisms that are not fully captured with data with a limited number of genetic markers (Emerson et al., 2010; Jeffries et al., 2016; Puckett et al., 2016; Marková et al., 2020). Using GBS data from over 500 individuals, the purpose of the present study was to reconstruct the phylogeographic history of the red fox in Europe by investigating broad-scale patterns of genomic variation, differentiation, and admixture amongst contemporary populations to investigate if contemporary red fox populations are the result of isolation in particular refugia during the LGM (Statham et al., 2018); or if these patterns likely emerged in the post-glacial period following the LGM (Edwards et al., 2012; Teacher, Thomas, & Barnes, 2011). From there, we adopted an Approximate Bayesian Computation (ABC; (Beaumont, Zhang, & Balding, 2002)) framework to distinguish between multiple post-glacial re-colonization scenarios of previously glaciated regions that were not fully resolved in previous studies using mtDNA, Y chromosomal, and microsatellite markers (Statham et al., 2018; Wallén et al., 2018). We first focused on the colonization of Ireland and Britain, and whether the red fox colonized the island of Ireland naturally or was introduced later by humans (Statham et al., 2018). Finally, we investigated the post-glacial colonization patterns of northern Europe, with a focus on the mechanism and timing of the colonization of Scandinavia around and after the Younger Dryas period (Wallén et al., 2018).

## Materials and Methods

### Laboratory methods

Red fox tissue samples were obtained from freshly culled (not directly related to this study), roadkill, frozen, ethanol-(70-95%) and DMSO-preserved material from previous studies. No material was collected for the purpose of the present study. Genomic DNA was extracted using the DNeasy Blood and Tissue Kit (Qiagen Ltd.) according to manufacturer’s protocols (with the additional treatment of RNase). A total of 30–100 ng/μl of high molecular weight DNA was sent to the Genomic Diversity Facility in Cornell University (USA) where GBS was used for constructing reduced representation libraries (Elshire et al., 2011) using the restriction enzyme EcoT22I (ATGCAT) in six GBS libraries (each consisting of 95 uniquely barcoded individuals and one negative control). Individual ligations were then pooled and purified from adaptor excess using the QIAquick PCR Purification Kit (Qiagen). For library preparation genomic adaptor-ligated fragments were then PCR amplified in 50 μL volumes with 10 μL of DNA fragment pool, 1 × Taq Master Mix (New England

Biolabs Inc.) and 12.5 pmol each of the following Illumina primers: 5′-<u>AATGATACGGCGACCACCGAGATCTACAC</u>TCTTTCCCTACACGACGCTCTTCCG ATCT and 5′-<u>CAAGCAGAAGACGGCATACGAGAT</u>CGGTCTCGGCATTCCTGCTGAACCGCTCT TCCGATCT (the underlined parts will hybridize to the two Illumina flowcell oligos). Temperature cycling consisted of 72 ◻C for 5 min, 98 ◻C for 30 s followed by 18 cycles of 98 ◻C for 30 s, 65 ◻C for 10 s, and 72 ◻C for 30 s, with a final extension step at 72 ◻C for 5 min. The EcoT22I GBS libraries (now containing ID tags and Illumina flowcell adaptors) were purified again using the QIAquick PCR Purification Kit (Qiagen). An aliquot was run on the BioAnalyzer™ 2100 to verify fragment sizes. Library DNA was then quantified on a Nanodrop 2000 (ThermoFisher Scientific) and subsequently sequenced on an Illumina HiSeq 2000 (Cornell University, Life Sciences Core Facility).

### Bioinformatics

The raw Illumina sequence data from 568 individuals were processed through the GBS analysis pipeline implemented in TASSEL v5.2.31 (Glaubitz et al., 2014). Due to concerns about the performances of *de novo* approaches to identify SNPs in reduced representation genomic techniques (particularly for demographic analyses; Shafer et al., 2017) the reference genome of the domestic dog (CanFam3.1; *Canis lupus familiaris*) was used to align the sequence tags on individual chromosomes using the BWA-backtrack method in the Burrows-Wheeler alignment tool (Li & Durbin, 2009).

A total of unique 9,249,177 tags were found, with 6,267,895 tags (67.8%) uniquely aligned to the dog genome. The rest of the tags were disregarded from further analyses. SNPs were initially filtered by removing those with a minor allele frequency (MAF) of <0.01 and missing data per site of >0.9. This resulted in 144,745 SNPs with a mean individual depth of 15.853 (±SD = 4.629). After the removal of indels, SNPs with an observed heterozygosity >0.5 (to filter out potential paralogs), removing SNPs and individuals with call-rates of <0.8, SNPs with a MAF of <0.05, and SNPs located on the X chromosome, a total of 19,795 bi-allelic SNPs and 524 individuals were retained after filtering in TASSEL v5.2.31 (Glaubitz et al., 2014) and PLINK v1.07 (Purcell et al., 2007). Because of the potential for linkage disequilibrium (LD) to bias the results of population genetic analyses, we pruned 3,571 SNPs with LD of r^2^ > 0.2 (Schweizer et al., 2016) in a window of 50 SNPs (sliding window with five SNPs overlapping at a time) in PLINK (Purcell et al., 2007). To filter out SNPs deviating from Hardy-Weinberg Equilibrium (HWE), individuals were grouped into 29 populations, consisting of seven or more individuals (see below). A least-squares F_IS_ estimator (based on an AMOVA) was implemented in GenoDive v2.0b23 (Meirmans & Van Tienderen, 2004) using 999 permutations. A total of 869 SNPs which deviated from HWE (*p* < 0.01) in 15 or more populations (≥ 51.7%) were removed from subsequent analyses (Van Wyngaarden et al., 2017).

Given that the focus of the study was to investigate demographic and phylogeographic patterns, loci putatively under selection were identified using Principal Component Analysis (PCA) and Mahalanobis distance implemented in the R package *pcadapt* v3.0 (Luu, Bazin, & Blum, 2017). A false discovery rate (FDR) of α = 0.05 was applied, with *q*-values smaller than α considered as candidate SNPs under selection. A total of 352 SNPs were removed according to these criteria. Two datasets were created for subsequent analyses; one containing all 524 individuals (*524dataset*; Table S1), and the other containing 494 individuals (*494dataset*; Tables 1 and S1), consisting of 29 ‘populations’ with seven or more individuals for population-level analyses (Fig. 1A-C).

**Table 1.** Population-level analyses of 29 pre-defined red fox populations. Populations are named by country of origin with an accompanying label (Pop ID) for illustrative purposes (see Figure 1). Approximate geographic co-ordinates are given in longitude and latitude. The number of individuals in each population is indicated by *N* and population-level measures of allelic richness (AR) and expected heterozygosity (He) are shown that were used to calculate the interpolation of genomic diversity (see Figure 3).

| Population | Pop ID | Longitude | Latitude | *N* | AR | He |
| --- | --- | --- | --- | --- | --- | --- |
| Ireland | IRE | 53.3927275 | -9.9262638 | 20 | 1.68 | 0.237 |
| Northern Ireland | NIR | 54.717149 | -6.219987 | 20 | 1.68 | 0.238 |
| England North | UKN | 54.47445462 | -2.680500725 | 17 | 1.73 | 0.254 |
| England South | UKS | 51.6839616 | 0.574959028 | 27 | 1.73 | 0.253 |
| Portugal North | PTN | 40.7115386 | -7.7495801 | 11 | 1.77 | 0.263 |
| Portugal South | PTS | 38.9648718 | -9.2992138 | 18 | 1.76 | 0.26 |
| Spain North | SPN | 42.829585 | -2.700338 | 29 | 1.79 | 0.268 |
| Spain South | SPS | 37.14280344 | -3.647460938 | 8 | 1.71 | 0.254 |
| France | FRA | 48.50605105 | 2.64715577 | 20 | 1.79 | 0.27 |
| Belgium | BEL | 50.589536 | 6.123152 | 19 | 1.81 | 0.274 |
| Switzerland | SWZ | 46.7633251 | 7.1564597 | 19 | 1.79 | 0.271 |
| Italy North | ITN | 45.926389 | 10.251667 | 7 | 1.74 | 0.269 |
| Italy | ITA | 43.32517768 | 11.31591797 | 20 | 1.73 | 0.254 |
| Germany | GER | 52.5200066 | 13.404954 | 28 | 1.81 | 0.273 |
| Slovenia | SOV | 46.31255967 | 15.50934385 | 24 | 1.78 | 0.265 |
| Croatia | CRO | 45.4082114 | 13.6589423 | 19 | 1.81 | 0.274 |
| Poland East | PLA | 53.82505665 | 23.3406601 | 16 | 1.82 | 0.274 |
| Poland South | PLS | 49.7801 | 20.2505 | 14 | 1.81 | 0.275 |
| Poland West | PLC | 52.117714 | 16.779255 | 19 | 1.82 | 0.275 |
| Serbia | SER | 45.83645405 | 20.43457031 | 18 | 1.81 | 0.271 |
| Ukraine | UKR | 47.725824 | 29.952977 | 9 | 1.8 | 0.274 |
| Sweden South | SWS | 57.95217685 | 13.93725559 | 17 | 1.74 | 0.256 |
| Sweden North | SWN | 64.3322949 | 20.28735358 | 16 | 1.77 | 0.263 |
| Norway | NOR | 70.26464 | 30.34619 | 14 | 1.77 | 0.265 |
| Finland North | FNN | 69.9059 | 27.02396 | 8 | 1.76 | 0.269 |
| Finland South | FNS | 60.46667 | 26.91667 | 12 | 1.76 | 0.259 |
| Russia Komi | RUK | 61.8 | 51.516667 | 10 | 1.76 | 0.262 |
| Russia Kirovskaja | RUV | 58.56861045 | 48.86787415 | 25 | 1.78 | 0.265 |
| Siberia | SIB | 135.774568 | 67.898166 | 10 | 1.6 | 0.211 |

**Figure 1.**
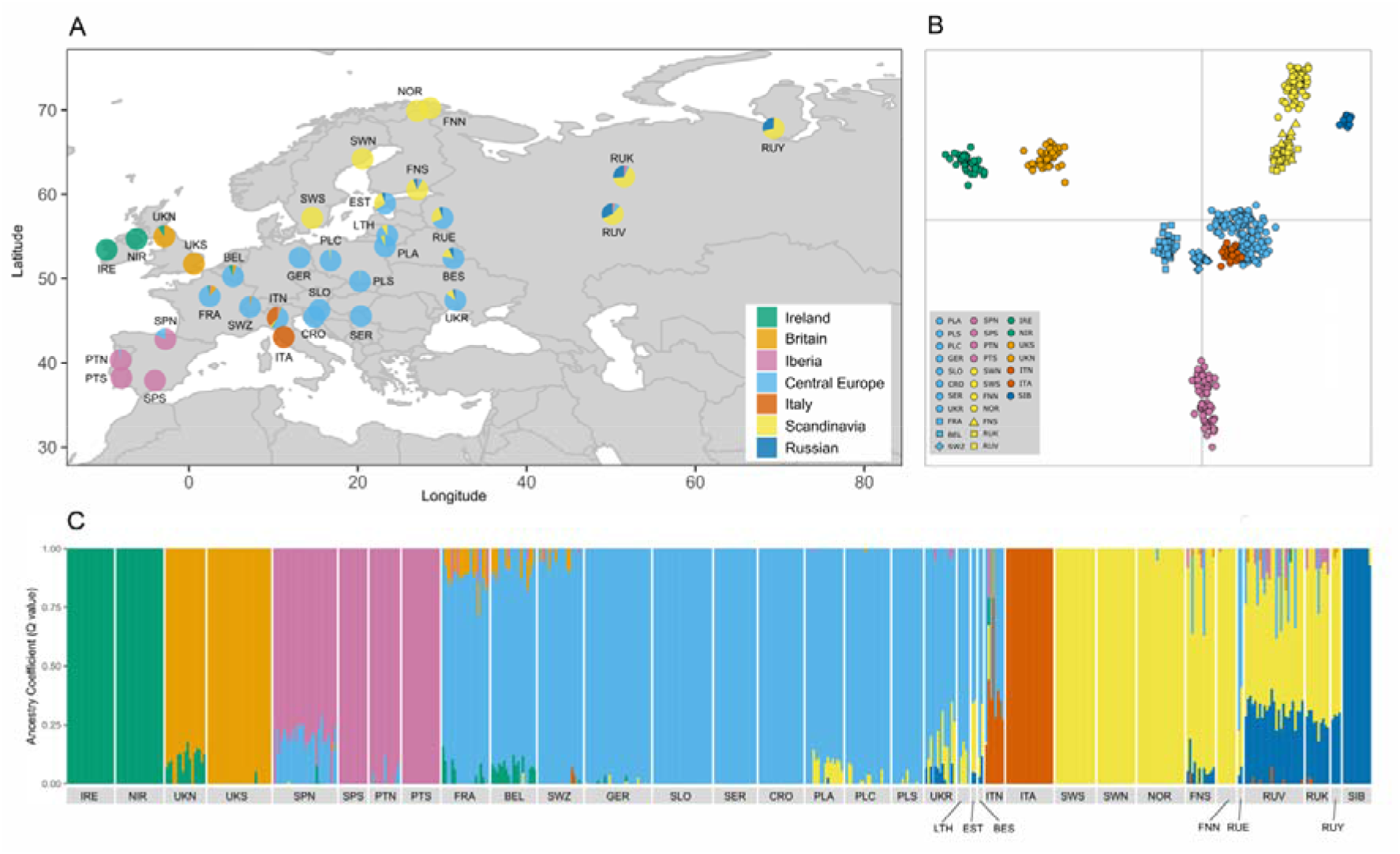
Approximate locations of the studied populations (A) and the genomic clusters to which they have been assigned based on Discriminant Analysis of Principal Components (DAPC; B) and Bayesian analysis in *fastSTRUCTURE* at *K* = 7 (C). The proportion of admixture in each population (A) is based on the ancestry coefficients determined in *fastSTRUCTURE* (C).

### Individual-based analyses

Individual-based clustering analysis on the *524dataset* was conducted in fastSTRUCTURE v1.0 (Raj, Stephens, & Pritchard, 2014). fastSTRUCTURE was performed using the simple prior with *K* values of 1–30 over five independent runs. The number of clusters (*K*) was obtained by using the ‘chooseK.py’ function on each of these independent runs. Visualization of individual assignments to clusters per population was initially performed using the ‘distruct.py’ function, with the final figure produced using a custom R script. In our second individual-based analysis on the *524dataset*, we incorporated the capture or sampling location of each individual, using the spatially-explicit software *TESS3* (Caye, Deist, Martins, Michel, & François, 2016) The default values of the program were implemented and each run was replicated five times. The optimal value of *K* corresponded to the minimum of the cross-entropy criterion, across values of *K* = 1–30.

### Population-based analyses

For the *494datset*, two measures of genomic diversity, allelic richness (AR) and expected heterozygosity (H_E_), were calculated in GENODIVE (Meirmans & Van Tienderen, 2004) and HP-RARE (Kalinowski, 2005). AR and H_E_ values were mapped with interpolation using ArcGIS 10.2.1. Geostatistical Analyst. One population from Siberia (Russia) was excluded from this analysis because it was geographically distant from all the other populations. Interpolation was carried out using an Inverse Distance Weight model (IDW, power=1, based on 12 neighbours; Stojak et al., 2016). Discriminant Analysis of Principal Components (DAPC) as implemented in the R package *adegenet* v2.0.1 (Jombart & Ahmed, 2011) was performed on the *494dataset*. DAPC does not make assumptions about Hardy-Weinberg equilibrium or linkage disequilibrium and provides a graphical representation of the divergence among pre-defined populations/groups.

### Phylogeographic reconstruction

Approximate Bayesian Computation was implemented in DIYABC Random Forest v1.0 (Collin et al., 2021) to further investigate the dynamics of the re-colonization process of red foxes in Europe. As the method is computationally intensive and has generally required 100,000 to 1,000,000 simulations to distinguish between demographic scenarios being tested, several studies have performed their ABC analyses on subsets of their SNPs randomly selected from the full dataset (usually ~1,000 SNPs; e.g. Huang et al., 2017; Jeffries et al., 2016). However, recent applications using a tree-based classification method known as ‘random forest’ in ABC allow demographic scenarios to be distinguished based on 1,000s to 10,000s of simulations for each scenario (Fraimout et al., 2017; Kotlík, Marková, Konczal, Babik, & Searle, 2018; Pudlo et al., 2015; Marková et al., 2020). We followed the approach of Kotlík et al., (2018) where all SNPs were used, but we chose a subset of the individuals in each grouping to save on computational time (Table S1).

Based on recent simulation studies, ABC-based methods have received criticism for their ability to capture the true demographic models under consideration (Cabrera & Palsboll, 2017; Shafer, Gattepaille, Stewart, & Wolf, 2015). Following recommendations by Cabrera & Palsboll (2017), scenarios were kept as simple and different from each other as possible in order to distinguish between the major demographic events under consideration. In addition, the number of comparable scenarios was always kept low (Cabrera & Palsboll, 2017). Following on from outstanding issues in regards to unresolved colonization scenarios (Statham et al., 2018), we first investigated the colonization history of red foxes in the British Isles. We grouped individuals into three ‘populations’ of interest to ascertain the most likely scenario for the timing and source of existing populations in Ireland and Great Britain based on the analyses performed in fastSTRUCTURE, DAPC (See Results) and previous studies. These were ‘Ireland’ (IRE + NIR populations), ‘Britain’ (UKS + UKN) and ‘Europe’ (FRA + BEL + SWZ). Three scenarios were incorporated that the data and analysis presented by Statham et al. (2018) could not distinguish between. The first is that Ireland was colonized naturally overland from an unsampled ancestral population (N4) after the LGM (i.e. before Ireland became an island between approximately 19,000 and 15,000 yrs BP; t2; Fig. S1). In this scenario, Britain originated from this ancestral population also but mixes with the European population before it became isolated from the mainland after the flooding of Doggerland around 8 kyrs BP (19–8 kyrs BP; ta). Europe diverged from the unsampled ancestral population at 19–15 kyrs BP with t1>t2. In the second scenario, Ireland’s red foxes were founded directly from Britain naturally before it became an island (t2). In the third scenario, Ireland’s foxes were transported by humans from Britain after the earliest evidence of human presence on the island at 12,700 yrs BP (Dowd & Carden, 2016) right up to the present day (t3; Fig. S1). Effective populations sizes were allowed to range between 10 and 500,000 individuals for Irish and British populations and between 10 and 1,000,000 for Europe and the unsampled ancestral population. Generation time was assumed to be 2 years (Statham et al., 2018).

For the second ABC-based analysis, we investigated the colonization history of Scandinavia. Wallen et al. (Wallén et al., 2018) proposed that Scandinavia was colonized from multiple sources based on mtDNA and Y chromosome data but did not attempt to date the progression of these events. We attempted to distinguish between five scenarios for the timing and progression of these colonization events. In the first of these, ‘Scandinavia’ (FNS + FNN + SWN + SWS + NOR) was the result of admixture in the east and subsequent colonization (9,000 yrs BP to present; tb) between ‘Central Europe’ (PLA + PLC + PLS + GER) and ‘Russia’ (RUK + RUV) populations after the region became ice-free after the Younger Dryas Glaciation and disappearance of land-bridges connecting it to central Europe (Herman et al., 2014). The Central Europe and Russian populations split at t1 (27,000–19,000 yrs BP). In the second scenario, the first colonization wave occurred from central Europe over land-bridges (14,000 to 9,000 yrs BP; ta), with later admixture occurring from Russia (9,000 yrs BP to present; tb). In the third scenario, the first colonization wave occurred from Russia (9,000 yrs BP to present; tb), with later admixture from Central Europe from an eastern route (9,000 yrs BP to present but restricted to after the first Russian colonization wave; tb >tc). In the fourth scenario, Russia was the result of admixture between Scandinavia and Central Europe at tb. Finally, the fifth scenario had all three populations diverging at t1. Effective populations sizes were allowed to range between 10 and 1,000,000 individuals.

Ten thousand simulated datasets per scenario were used to produce posterior distributions. Each scenario was considered to be equally probable at the outset. To check the reliability of the observed summary statistics, a Principal Component Analysis (PCA) was performed on the summary statistics from the simulated datasets and compared against the summary statistics from the observed dataset in order to evaluate how the latter is surrounded by the simulated datasets (Collin et al., 2021). All simulated datasets were used in each Random Forest training set. We used five noise variables and ran 1,000 Random Forest trees to select the most likely scenario and estimate parameters (Collin et al., 2021).

## Results

### Individual-based analyses

fastSTRUCTURE identified *K* = 7 as the lower limit of clusters in each of the five independent runs of *K* = 1–30 (Fig. 1C), with the upper limit of *K* fluctuating from 10-13 between runs. Focusing first on *K* = 7, distinct clusters were identified in each of Ireland (’Ireland’) and Great Britain (’Britain’). Iberian populations formed a distinct cluster (’Iberia’), a known glacial refugium. Populations in France, Switzerland, Belgium, Germany, Poland, Slovenia, Croatia, Serbia and the Ukraine formed a distinct cluster (’Central Europe’, named for simplicity because of the approximate location of the cluster relative to the other clusters). Several of these populations are in proximity to the Balkan and Carpathian glacial refugia. Localities with small numbers of individuals in Lithuania, Estonia, Belarus and western Russia also belonged to this cluster. These populations in eastern Europe showed evidence of mixed ancestry with individuals from Scandinavia (Figs. 1A and 1C), who formed another distinct cluster (‘Scandinavia’). Individuals from European Russia had mixed ancestry between this Scandinavian cluster and individuals from Siberia (another distinct cluster; ‘Siberia’). Finally, individuals from central Italy formed a distinct cluster (‘Italy’), another well-defined glacial refugium. Individuals in northern Italy showing mixed ancestry with the central European cluster (Figs. 1A and 1C). For the lower values of *K*, the islands of Ireland and Britain were separated from all other samples at *K* = 2. Scandinavia and Russia samples were further separated at *K* = 3, Iberia samples at *K* = 4, Siberia at *K* = 5 and Ireland and Britain each becoming distinct clusters at *K* = 6 (Fig. 2). For *K* = 8, an additional cluster with the European Russian populations was identified. For *K* = 9, another cluster consisting of French, Belgian and Swiss populations was found (with mixed ancestry from Central Europe; Fig. 2). These populations are in close proximity to the known glacial refugium located in the Dordogne region. Further admixture was identified within populations in Central Europe at *K* = 10–13 (data not shown). The spatially-explicit software *TESS3* (Caye et al., 2016) failed to resolve the genomic structure within the system, with the minimum of the cross-entropy criterion decreasing up to *K* = 30 (Fig. S4). Further test runs increasing *K* up to a maximum of 40 failed to resolve the genomic structure (data not shown).

**Figure 2.**
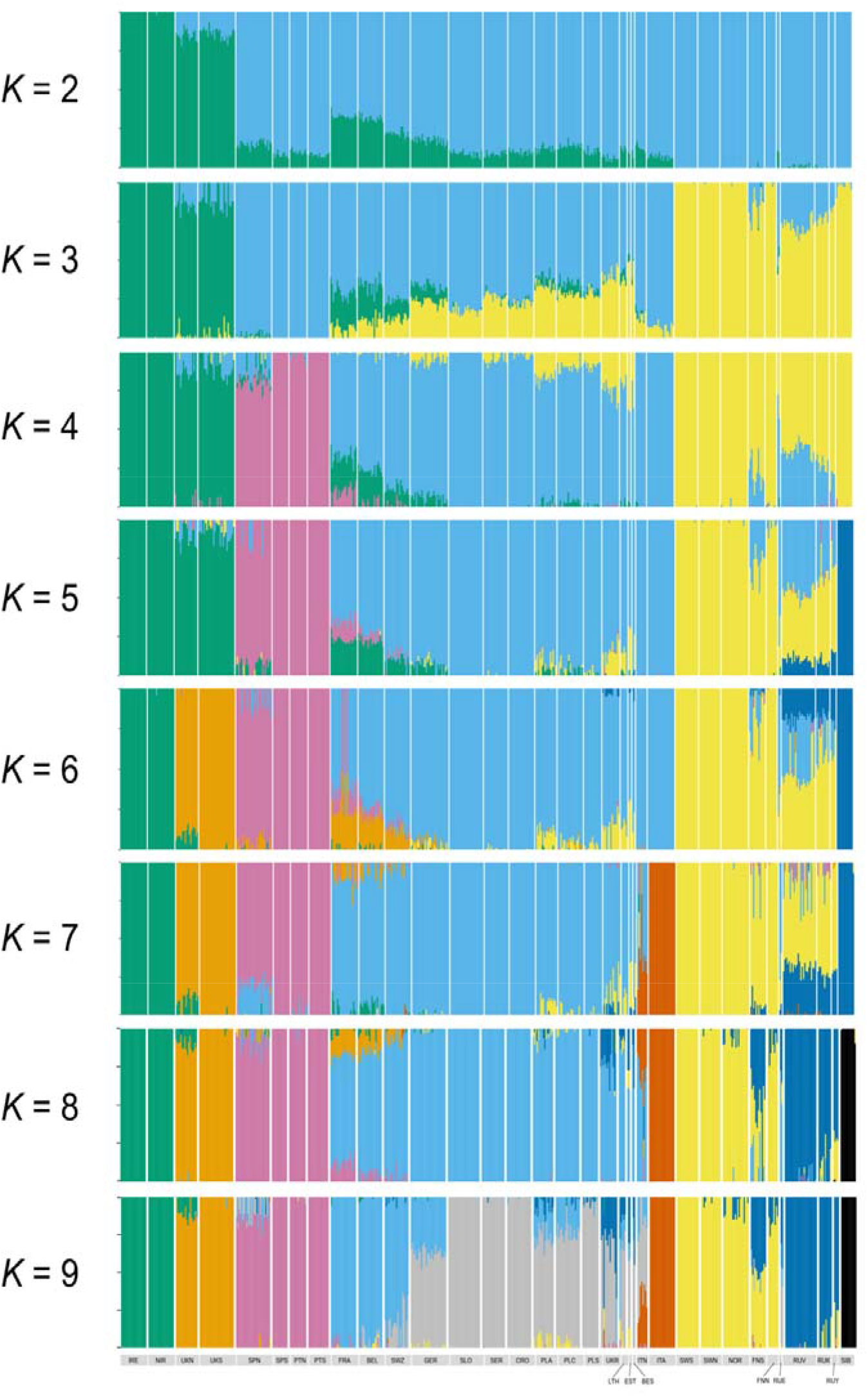
The proportion of admixture in each population based on the ancestry coefficients determined in *fastSTRUCTURE* for *K* values 2–9.

### Population-based analyses

Direct estimates of genomic diversity (Table 1) and the IDW interpolation of allelic richness (AR) and expected heterozygosity (H_E_) in 28 fox populations showed that diversity is highest in Central and Eastern Europe and decreases westwards and northwards (Fig. 2). Genomic diversity was notably lower in the British Isles, with the Irish populations showing the lowest levels of diversity (Fig. 2). The DAPC revealed distinct groupings of Iberian samples, Irish samples, British samples, Siberian and Scandinavia/Russian samples in general agreement with the individual-based analysis in fastSTRUCTURE (Fig. 1B). Populations in western, central and eastern Europe were grouped closely together, but the populations in France, Belgium and Switzerland were more separated from the main European group on the first axis, aligned with the individual-based Bayesian analyses at *K* = 9 (Fig. 2). Although the population in central Italy formed its own genomic cluster in the individual-based analysis (Fig. 1C), it grouped more closely with the central European group than the French, Belgian and Swiss populations in this analysis (Fig. 1B).

For the ABC-based analysis, first focusing on the colonization history of Ireland and Britain, Scenario 1 (Fig. S1) was selected (620/1,000 trees) with the posterior probability estimated at 0.741. The ancestral population for Ireland and Britain was estimated to have split from the European mainland at 17.7 kyrs BP (95% CI: 15.81–18.95 kyrs BP), with the Irish population originating at 16.1 kyrs BP (95% CI: 15.04–18.1 kyrs BP; Fig. 4; Table S2). For Scandinavia, Scenario 1 (510/1000 trees) was selected with the posterior probability estimated at 0.66. The initial split of the Central European and Russian populations was estimated at 23.04 kyrs BP (95% CI: 19.34–26.55 kyrs BP). The current Scandinavian originates from admixture from Russian and Central European populations at 5.36 kyrs BP (95% CI: 1.83–8.75 kyrs BP; Fig. 4; Table S2).

## Discussion

In this study, we provided a genome-wide assessment of population structure and diversity in the red fox in Europe. By incorporating over 15,000 SNPs and over 500 individuals, we were able to advance previous work by investigating broad-scale patterns of structure and variation to identify putative glacial refugia and post-glacial re-colonization patterns in this widespread species.

### Phylogeographic structure of the red fox in Europe

Individual- and population-based analyses revealed congruent patterns of genomic structuring at the broad scale of Europe (and Siberia), with certain important nuances being revealed by different approaches (Figs. 1, 2, 3 and S4). Earlier studies had proposed that red foxes may have existed as a single, large panmictic population during the LGM based on a lack of distinct structure at mtDNA markers using modern and/or ancient DNA (Edwards et al., 2012; Teacher et al., 2011). If this was the case, we might have expected to find a more continuous population (excluding the islands potentially) and this is not evidenced here with this greatly expanded dataset in terms of genetic markers, individuals and spatial coverage. In addition, a continuous population over the whole continent at the LGM is not generally congruent with the fossil data and the lack of fossil records beyond the more accepted refugial regions (Sommer & Benecke, 2005; Sommer & Nadachowski, 2006). Our study demonstrates that the observed patterns of genomic variation in contemporary red fox in Europe were mainly shaped by distinct refugial populations, with subsequent post-glacial admixture and isolation when this species had expanded into what is now its current distribution range in Europe (Sommer & Benecke, 2005).

**Figure 3.**
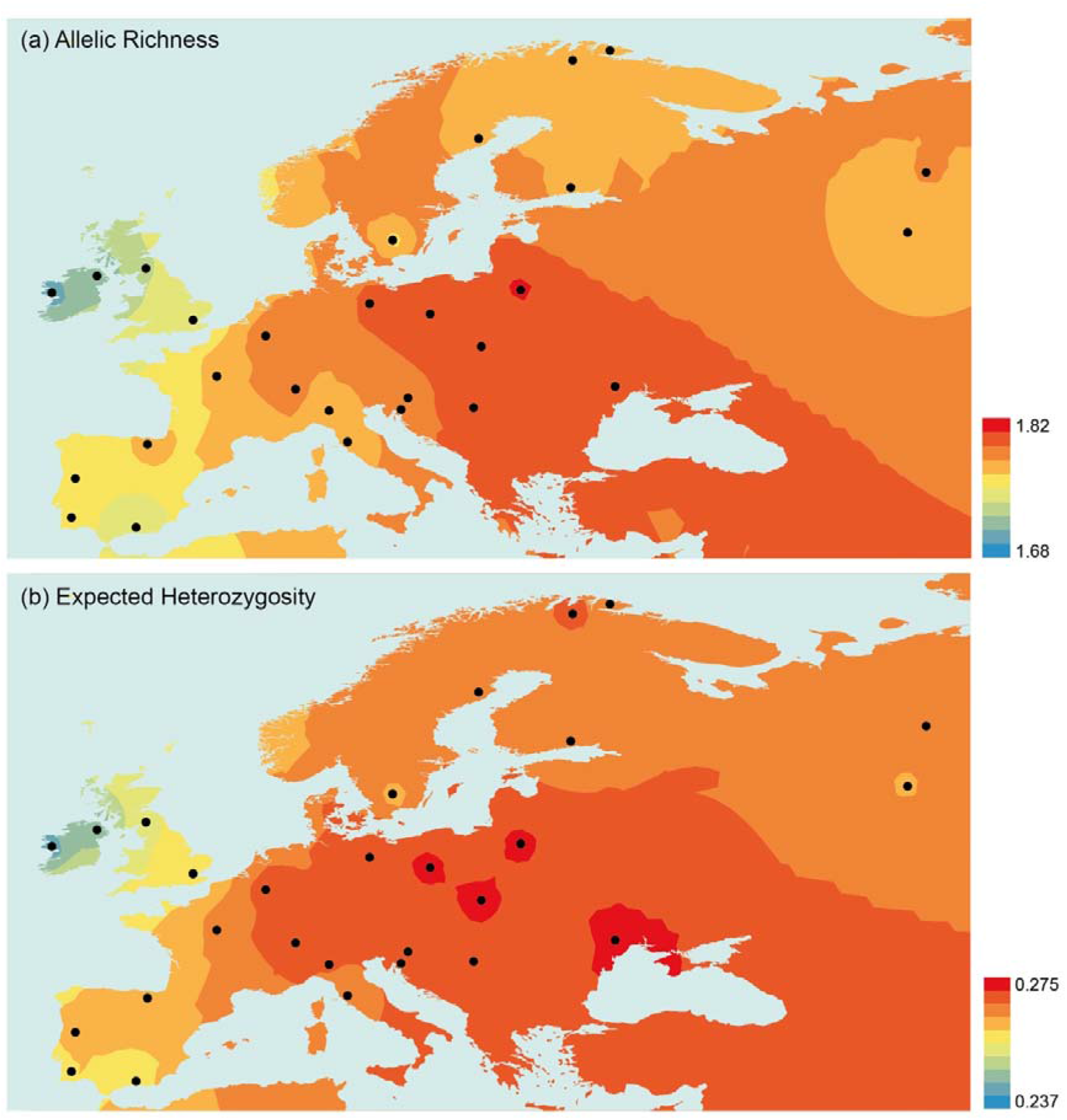
Interpolation of allelic richness and expected heterozygosity in 28 red fox populations (Siberia was excluded in this analysis) in Europe. The interpolated values of both indices are presented in the maps in different colours on a low (blue) to high (red) scale according to the legends. Black circles indicate the population locations.

**Figure 4.**
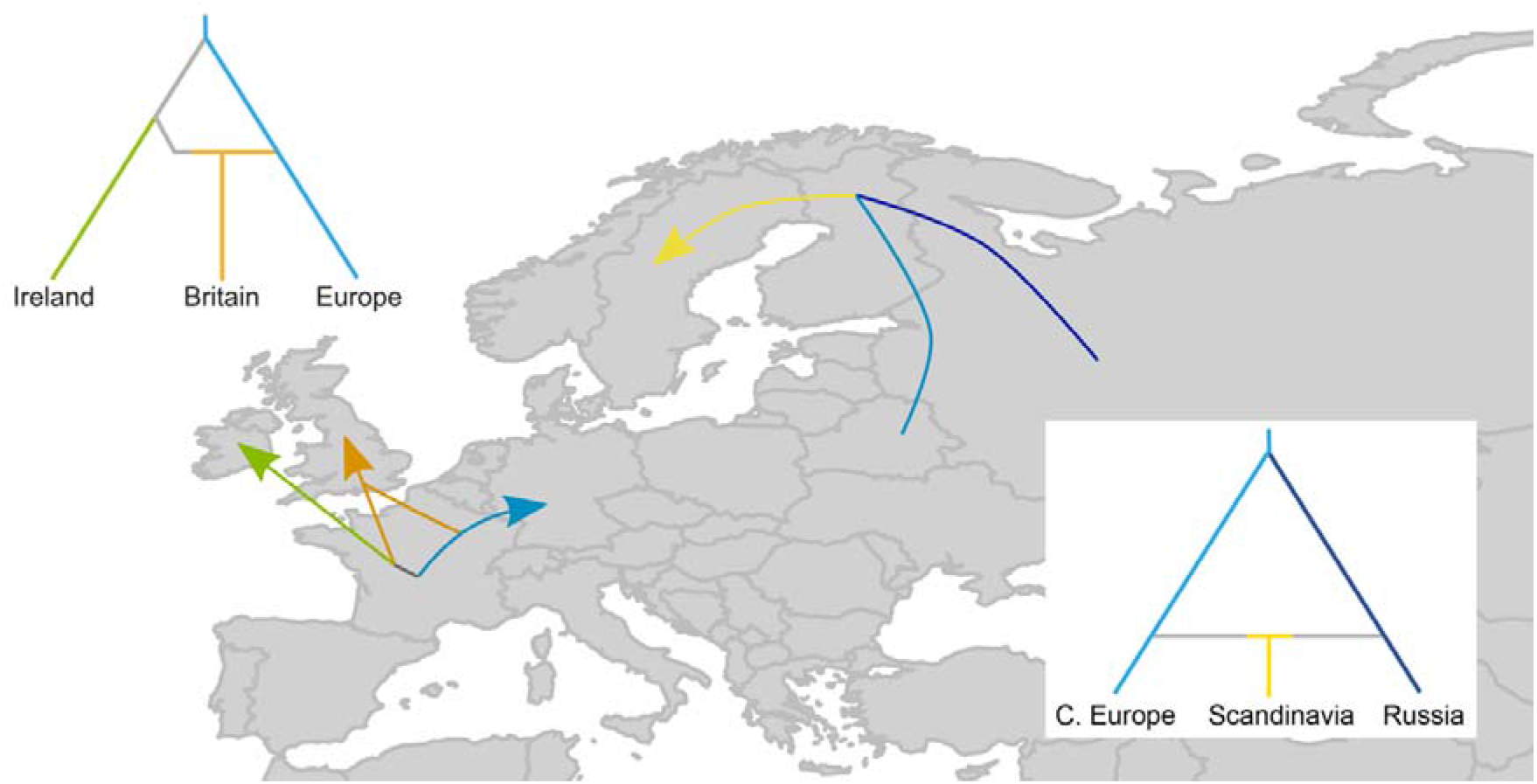
Graphical representation of the most likely post-glacial colonization scenarios for Ireland and Britain (A) and Scandinavia (B) inferred from Approximate Bayesian Computation.

At lower values of *K*, the islands of Ireland and Britain are separated from the rest of the European continent. Scandinavia/Russia and Iberia are then separated from the other continental individuals at *K* = 3 and 4, respectively (Fig. 2). Most of the central European (defined here as those outside of the Mediterranean peninsulas and Scandinavia) and Balkan populations formed a single genomic cluster at *K* = 7 (Figs. 1A and 1C). This and the elevated values of genomic diversity (Fig. 3) are likely reflective of more widespread and connected populations occupying the Balkans and Carpathian refugia during the LGM (as is known from fossil records; Sommer & Nadachowski, 2006) and a subsequent expansion into the rest of central Europe in the post-glacial period. A similar scenario has been proposed for other large mammals (Frantz et al., 2014; Stojak & Tarnowska, 2019). At *K* = 7, French, Belgian and Swiss individuals were grouped with other central European populations but population-level analyses (DAPC; Fig. 1B and 3) showed that these populations were distinct from other populations in close proximity (and they formed their own cluster at *K* = 9 in the individual-based Bayesian analysis; Fig. 2). Fossil records of the red fox are known from the Dordogne region in France during the LGM (Sommer & Nadachowski, 2006) so these populations may stem from a previously isolated refugial population in the area, and now show post-glacial admixture with populations stemming from eastern/Balkan and Iberian refugia (Figs. 1B and 1C). The Iberian populations form a distinct cluster/group at *K* = 4 and above (Figs. 1B, 1C and 2) and this is in line with previous findings using fewer molecular markers identifying this as a glacial refugium. Statham et al. (2018) identified mtDNA haplotypes that were endemic to the region, while microsatellites identified Spanish individuals as being distinct from those in other European populations. A similar pattern was found previously in badgers, with Iberian populations having many unique mtDNA haplotypes not found elsewhere on the continent (Frantz et al., 2014). The Pyrenees Mountains have remained a formidable barrier for post-glacial re-colonization, and there appears to be little contribution to subsequent northwards expansion when the ice-sheets receded for many terrestrial species (Bilton et al., 1998). Even though the maximum dispersal capabilities of the red fox are up to 1,000 km in Europe (Walton, Samelius, Odden, & Willebrand, 2018), this mirrors the pattern of mountains acting as significant barriers for the species in North America (Sacks, Statham, Perrine, Wisely, & Aubry, 2010). This is in contrast to red foxes in the other Mediterranean refugium, Apennine (Italy). Although red foxes from central Italy are identified as a distinct cluster in fastSTRUCTURE at *K* = 7 and above, mixed ancestry was identified with neighbouring populations north of the Alps in central Europe and the Balkans (Figs. 1B, 1C and 2). This may rather reflect a more recent divergence of this population given the geography of the region.

Glaciated regions during the LGM such as the British Isles and Scandinavia present differing problems in terms of how contemporary populations of terrestrial species colonized these areas in post-glacial periods. Ireland has existed as an island for approximately 15,000 years (Edwards & Brooks, 2008) and humans have been proposed as the primary mechanism of transport for its mammalian fauna based on existing fossil data and numerous phylogeographic studies (e.g. McDevitt et al., 2011; Carden et al., 2012; Frantz et al., 2014). Statham et al. (2018) proposed a split of approximately 10 kyrs between Irish and British foxes but with confidence intervals that didn’t distinguish between natural and human-mediated colonization scenarios. Both Irish and British populations are distinct from mainland European populations (Figs. 1B, 1C and 2) and have patterns of diversity and structure consistent with colonization and subsequent isolation (Table 1; Figs. 1B and 2). Using an ABC-based approach, a scenario in which Ireland was colonized before humans were known to be present (approximately 15 kyrs BP) was deemed the most likely (Fig. 4; Table S3). Although this conflicts with the current fossil evidence where the oldest known specimen in Ireland is from the Bronze Age (Sommer & Benecke, 2005), this is congruent with previous studies demonstrating high haplotype diversity and the identification of many unique haplotypes at mitochondrial markers on the island in comparison to British and other European populations (Edwards et al., 2012; Statham et al., 2018). The stoat was also proposed to be an early colonizer of Ireland over a post-glacial land bridge (Martínková, McDonald, & Searle, 2007) and several potential prey species (e.g. mountain hare *Lepus timidus* and arctic lemming *Dicrostonyx torquatus*; Woodman et al., 1997) were also present in the early post-glacial period. A recent re-assessment of several Irish mammalian fossils has pushed several species (reindeer *Rangifer tarandus*, grey wolf *Canis lupus* and woolly mammoth *Mammathus primigenius*) to the LGM/post-glacial boundary (Carden et al., 2020). Given that models of glaciation/de-glaciation patterns are reliant on secure and accurate fossil data, the island may have hosted a larger mammalian community in the early post-glacial period and this may have included the red fox also. While humans were still likely an important factor in determining later faunal assemblages on Ireland (Carden et al., 2012; Frantz et al., 2014), the early post-glacial period clearly warrants further investigation on the island based on the results presented here and newly available fossil data (Dowd & Carden, 2016; Carden et al., 2020).

Although glacial refugia in Scandinavia during the LGM have been proposed for several species including mammals (Lagerholm et al., 2014; Westergaard et al., 2019), the red fox does not appear in the fossil records in southern Scandinavia until much later (~9,000 yrs BP; Sommer & Benecke, 2005). Here, we examined five different hypotheses for the colonization of Scandinavia (Fig. S2). The most likely scenario for the colonization of Scandinavia is a mixture of foxes from Russian and central/eastern Europe colonizing from the east (Fig. 4). When the ice retreated from northern Scandinavia after the Younger Dryas, a lack of geographic barriers led to later dispersal into the region from the east (Norén et al., 2015; Wallén et al., 2018), a pattern that is evident in other carnivores also (Dufresnes et al., 2018; Keis et al., 2013). Several recent studies have pointed to a pattern of post-glacial colonization of Scandinavia from the south involving individuals from present-day central Europe first crossing temporary land-bridges prior to the Younger Dryas glaciation. This pattern is particularly prevalent in small mammals such as the field vole *Microtus agrestis* (Herman et al., 2014) and the bank vole *Myodes glareolus* (Marková et al. 2020). From all the analyses presented here, this scenario is not supported for the red fox and it is instead apparent that this came later from the east only (Fig. 4). With Scandinavia being one of the last regions of the continent to be re-colonized, the higher genomic diversity observed here than some of the more westerly regions could be due to it being populated from multiple sources from the east (Wallen et al., 2018; Marková et al., 2020).

Using genome-wide data has allowed us to tease apart broad-scale patterns of structure and diversity in a widespread carnivore in Europe that was not evident from more limited marker sets. The use of genomic data allowed us to identify the importance of refugial regions in terms of both endemism (e.g. Iberia) and sources of post-glacial re-expansion across the continent (e.g. the Carpathians and Balkans). In conjunction with ABC-based analyses, we identified patterns of post-glacial colonization in formerly glaciated regions that contradict previously proposed routes for the red fox and other similar species. Given the genomic resources now available (Kukekova et al., 2018), the application of ancient genomics on the extensive fossil material available for this species (Sommer & Benecke, 2005) should fall into line with other charismatic carnivores (Loog et al., 2020) to fully understand re-colonization and temporal patterns that have not been captured in previous studies of ancient red fox specimens (Edwards et al., 2012; Teacher et al., 2011).

## Acknowledgements

This study was financed by the National Science Centre, Poland, grant no. DEC-2012/05/B/NZ8/00976 awarded to JMW, ADM, RK, ER and CP. We thank the following people for supplying samples: Heikki Ahola, Peter Allason, Evidio Bartolini, Lucia Burrini, Benoit Combes, Dorothee Ehric, Teresa García Díez, Vaclavas Gedminas, Christian Gortázar Schmidt, Rebecca Hari, U.A. Kalisnikov, Marta Kołodziej-Sobocińska, Nikolay Korablev, Antonio Lavazza, Sandro Lovari, Luciano Palazzi, Giorgia Romeo, Marie-Pierre Ryser-Degiorgis, Ivan Seryodkin, Aleksandr Sokolov and Pavel Veligurov. ADM thanks Robert Sommer, Norbert Benecke and Ruth Carden for information on, and access to, their red fox fossil data, and to Petr Kotlik for advice on ABC analyses. We sincerely thank the three anonymous reviewers for their comments which significantly improved the manuscript.

## Author contributions

JMW, ADM, RK, JM, CP and ER conceived and acquired funding for the study. ADM, JMW, RK, MJS, BNS, CP and ER designed the study. MJS, ACF, KN, EÅ, JL, MB, CF, PS, DGT, MS, AG, MI, MP, AK, IMO, AS, BP, KF, VL, SWRH, DC, ER, BNS and RK contributed samples and data towards the study. IR performed the laboratory work. ADM, AR-G and IC generated the final SNP panel. ADM, IC, SSB, LR and JMW designed and performed the data analyses. ADM, JMW, IC, JS and SSB wrote the manuscript, with all authors contributing to edits and discussion.

## Data Accessibility

Raw SNP data and datasets used for individual and population-level analyses are available on a private peer review link on Dryad (https://datadryad.org/stash/share/lxGNZUG315NeDDYqLZ4srLJGzEqw3U7LLfLztMpaI6E)

## Notes

### Competing Interest Statement

The authors have declared no competing interest.

### Summary of Updates

Updated ABC-based analyses, leading to differences in the colonizaton sceanrio inferred for Scandinavia. More detailed STRUCTURE-based analyses also presented.

