## Supplementary Material for "Next-generation phylogeography resolves post-glacial colonization patterns in a widespread carnivore, the red fox (*Vulpes vulpes*), in Europe"

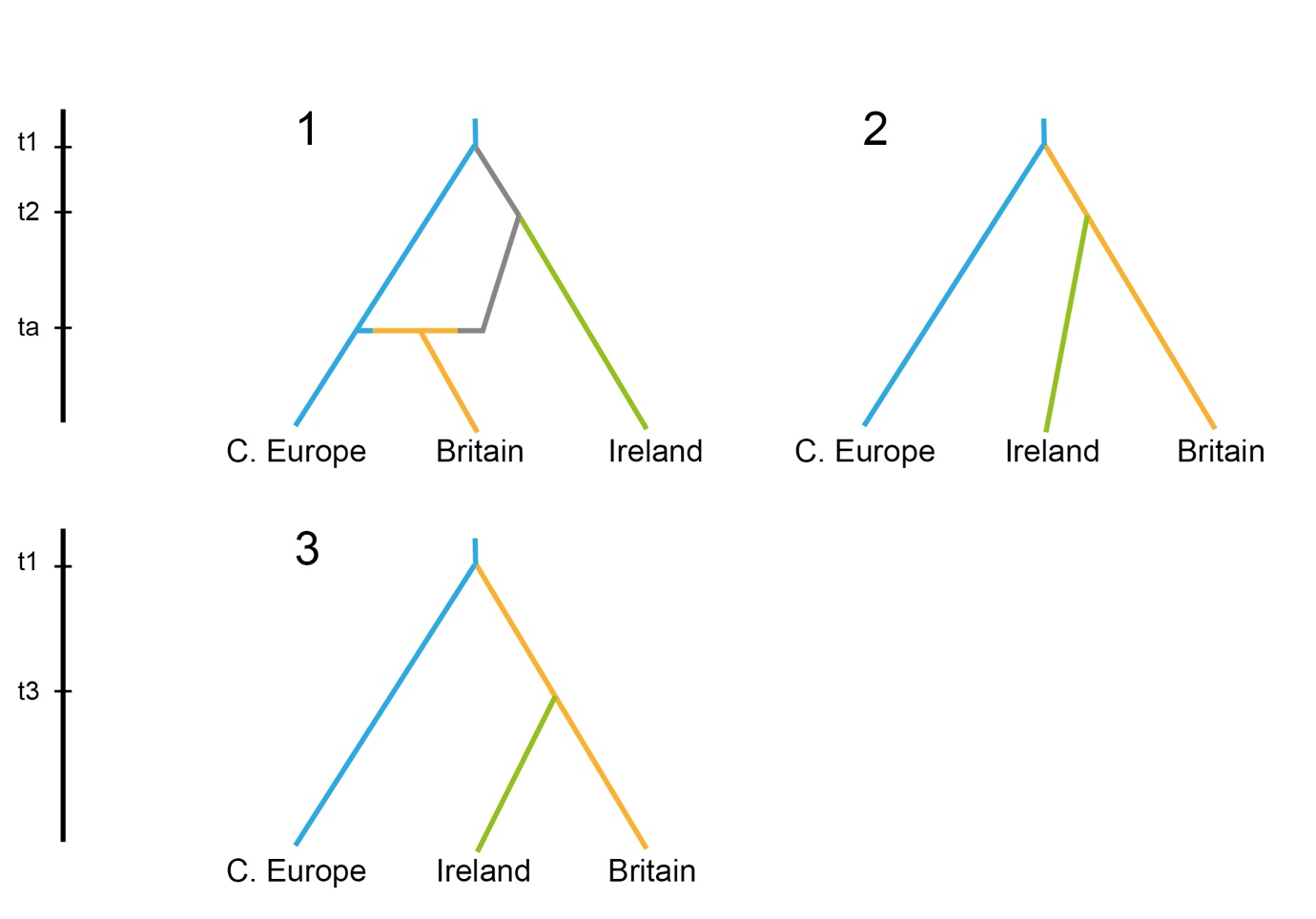

**Figure S1.** Colonization scenarios for the British Isles with Approximate Bayesian Computation and random forest. Effective population sizes were allowed to vary

between extant as well as ancestral lineages, as denoted by the different colours. See the main text for a more detailed description of the scenarios.

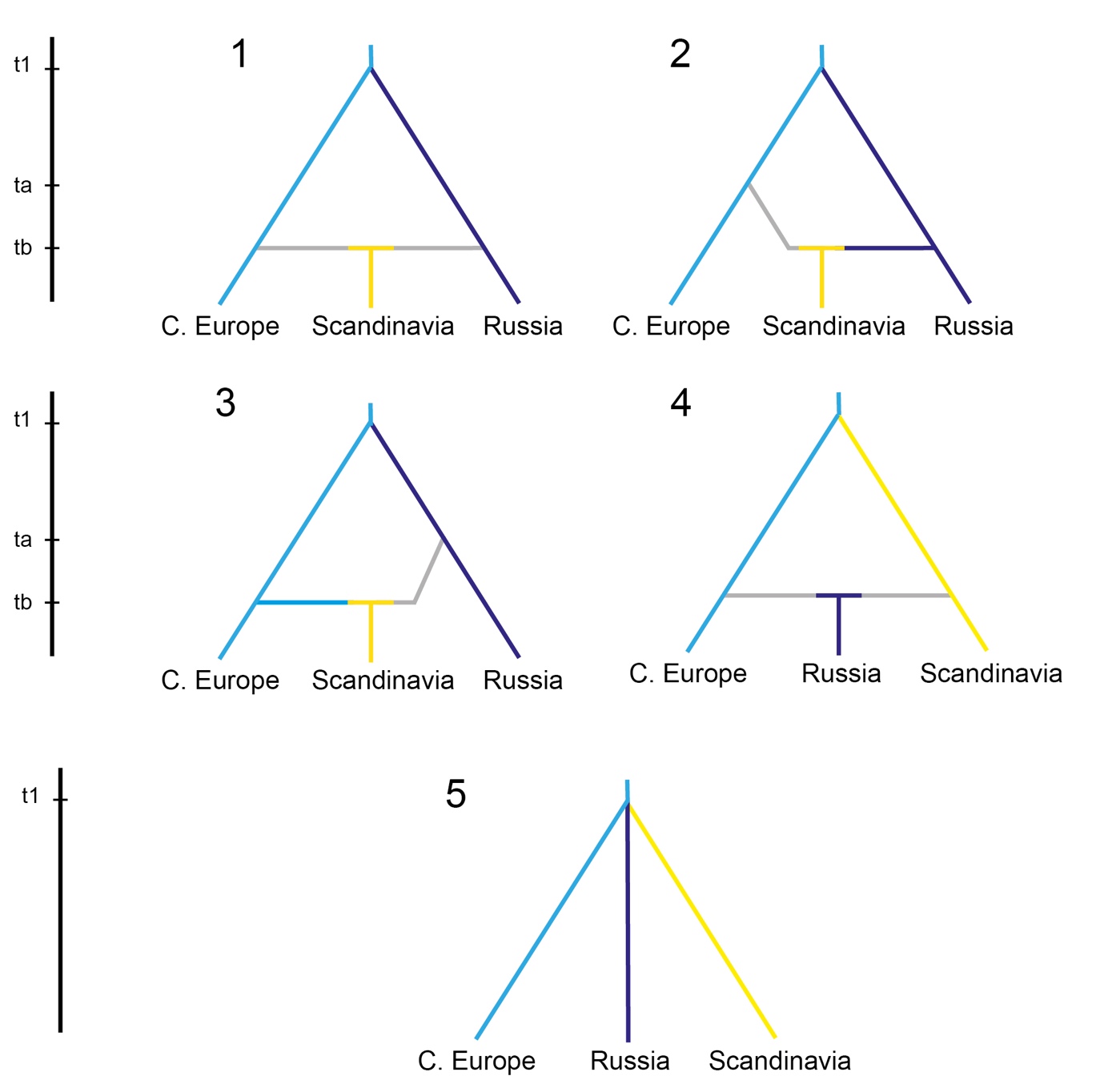

**Figure S2.** Colonization scenarios for Scandinavia with Approximate Bayesian Computation and random forest. Effective population sizes were allowed to vary

between extant as well as ancestral lineages, as denoted by the different colours. See the main text for a more detailed description of the scenarios.

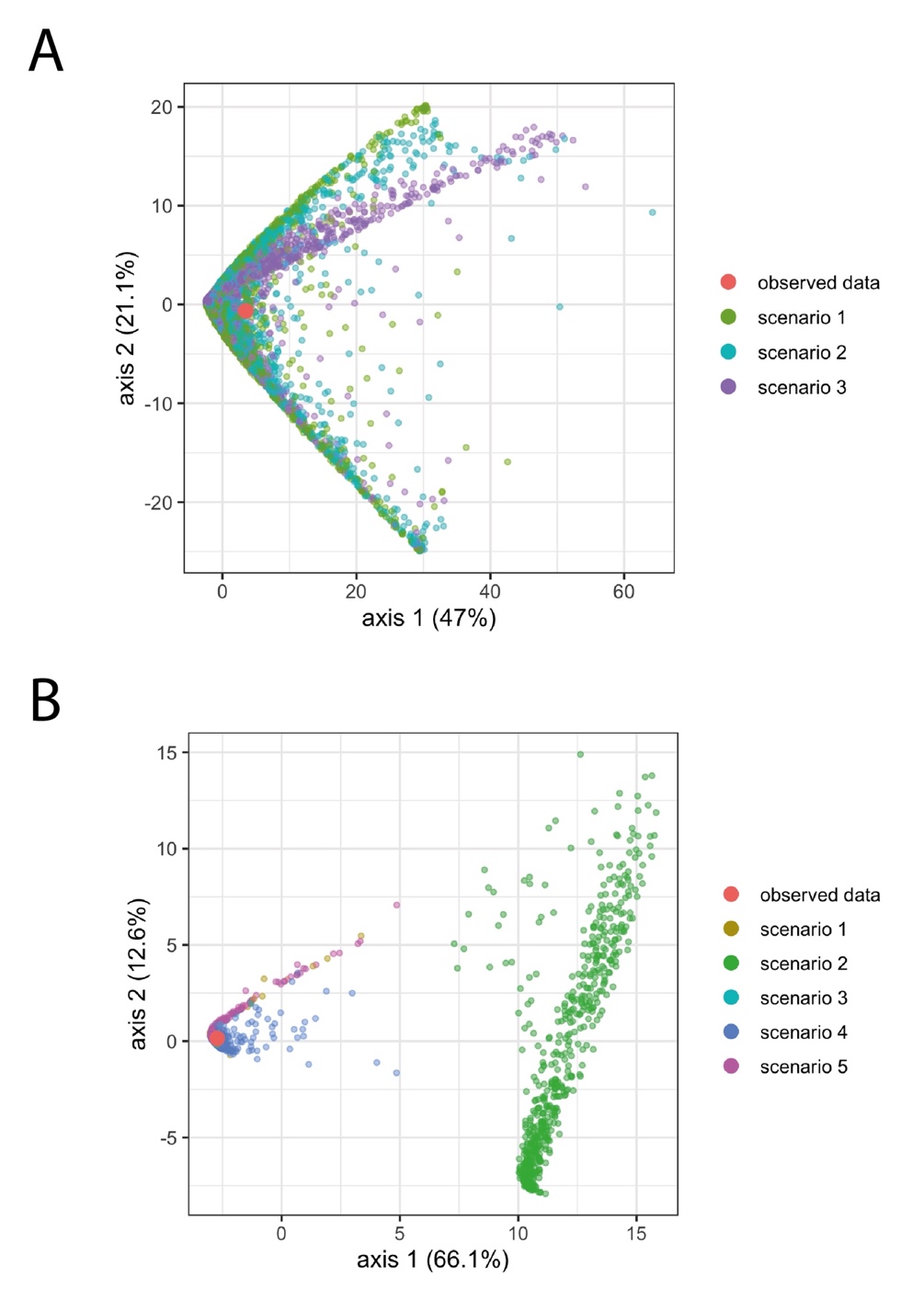

**Figure S3.** The fit of the data used in the ABC analysis of population history of red foxes are visualized using principal components analysis (PCA), in which the observations are the simulated data sets and the variables are the summary statistics. This shows the fit of the dataset (‘Observed data set’) in comparison to 10,000 simulated datasets of the scenarios for the colonization of the British Isles (A; see Fig. S1) and Scandinavia (B; see Fig. S2).

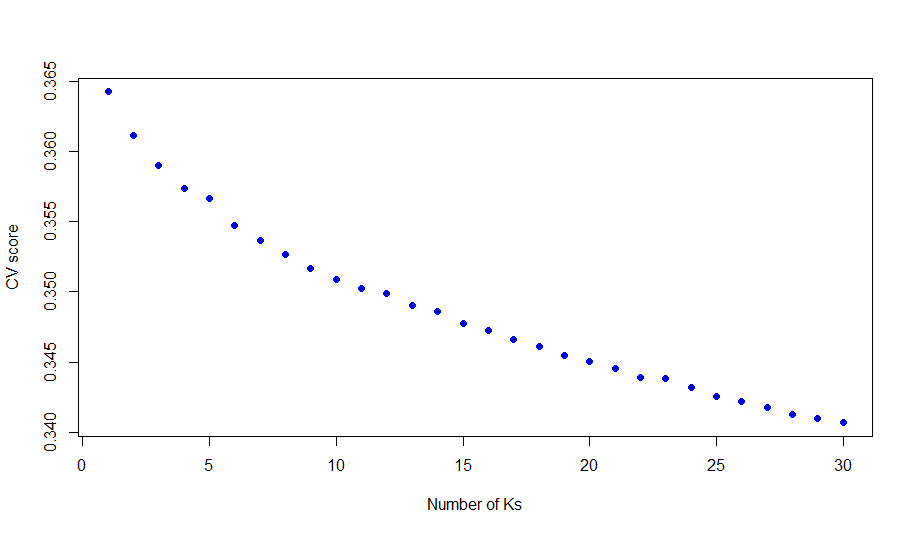

**Figure S4**. Cross-validation scores of *K* values 1-30 in *TESS3* (Caye et al., 2016)

**Table S1.** Details of 524 red fox samples used in the final analyses. These include the ‘population’ and individual IDs, the country, region and approximate locality of origin with corresponding latitude and longitude co-ordinates. Samples included in population-level analyses and Approximate Bayesian Computation (ABC) scenarios are indicated by an ‘X’ in the designated column. Table S1 provided as a separate .xlsx file.

**Table S2.** Parameter estimates for Scenario 1 (Colonization of British Isles) and Scenario 2 (Colonization of Scandinavia) and associated 95% Confidence Intervals defined by the 0.025 and 0.975 quantiles (*Q*) of the posterior distribution. Units are number of individuals for effective population size parameters (*N*) and years before present (yrs BP) for divergence time parameters (t).

| **Colonization of British Isles** |  |  |  |
| --- | --- | --- | --- |
| **Parameter** | **Median** | **Q0.025** | **Q0.975** |
| Ne (Ireland) | 301010 | 64177 | 480840 |
| Ne (Britain) | 245093 | 50964 | 480210 |
| Ne (Europe) | 769272 | 342570 | 979946 |
| t1 (yrs BP) | 17690 | 15808 | 18950 |
| t2 (yrs BP) | 16096 | 15038 | 18096 |
| ta (yrs BP) | 11980 | 8286 | 14712 |
| ra | 0.497 | 0.045 | 0.917 |
| **Colonization of Scandinavia** |  |  |  |
| **Parameter** | **Median** | **Q0.025** | **Q0.975** |
| Ne (Scandinavia) | 37159 | 10259 | 121373 |
| Ne (Russia) | 236058 | 58679 | 942023 |
| Ne (C. Europe) | 169136 | 73909 | 921580 |
| t1 (yrs BP) | 23042 | 19338 | 26550 |
| tb (yrs BP) | 5360 | 1832 | 8750 |
| ra | 0.587 | 0.379 | 0.803 |
